# Tonkean macaques do not prefer the helper or the hinderer in the hill paradigm

**DOI:** 10.1101/2025.02.28.640767

**Authors:** Marie Hirel, Hélène Meunier, Hannes Rakoczy, Julia Fischer, Stefanie Keupp

## Abstract

Evaluating others’ prosocial tendencies can benefit individuals by allowing them to interact with prosocial individuals and avoid antisocial ones. The ontogeny of humans’ strong prosocial preference has been widely investigated using the hill paradigm. Children’s preference for helper over hinderer agents was measured after they watched a scene in which the helper agent pushed a climber up a hill while the hinderer agent pushed the climber down the hill. Bonobos tested with the hill paradigm preferred the hinderer over the helper, contrasting previous findings for other nonhuman primates. We tested Tonkean macaques (*Macaca tonkeana*) using the same procedure as the one used with bonobos to see whether they would also exhibit a hinderer preference. Subjects did not prefer the helper over the hinderer (or vice versa). The low attentional level observed in our subjects suggests a lack of interest in the video stimuli. This finding relates to more general questions regarding how animals perceive abstract animated onscreen stimuli and the relevance of the hill paradigm in investigating prosocial preferences. Studies using various experimental paradigms with conspecifics or human actors as social agents are needed to investigate further the social evaluation of prosocial behaviours in Tonkean macaques, bonobos, and other primates.

## Introduction

Monitoring others’ prosocial behaviours (*i.e.*, actions that benefit others at a cost to oneself) is beneficial for animals to assess which potential social partners are likely to provide benefits to them (Barclay, 2016; Schino & Aureli, 2009; Schweinfurth & Call, 2019). Humans are susceptible to others’ prosocial acts and develop strong sociomoral preferences, *i.e.*, preferring cooperators and helpers while avoiding or even punishing uncooperative and non-helper individuals (Feinberg et al., 2014; Heiphetz & Young, 2014; Holvoet et al., 2016; Lyle & Smith, 2014). Nonhuman animals tend to reciprocate prosocial behaviours to individuals who have helped in the past (Dugatkin, 1997; Jaeggi & Gurven, 2013; Schweinfurth & Call, 2019; see also Clutton-Brock, 2009). Nonhuman primates (Anderson et al., 2013, 2016; Brügger et al., 2021; Herrmann et al., 2013; Kawai et al., 2014; Russell et al., 2008; Subiaul et al., 2008) and dogs (Chijiiwa et al., 2015; Kundey et al., 2011; Marshall-Pescini et al., 2011; Nitzschner et al., 2012) even demonstrated to discriminate and prefer prosocial over antisocial human experimenters.

The ontogeny of human prosocial preference has been mainly investigated in developmental psychology using the hill paradigm developed by Hamlin et al. (2007). Children could observe several times a scene with abstract social agents (geometrical shapes or puppets). In the helper condition, the scene starts with a character (the climber) repeatedly attempting to climb a hill without success. A second character (the helper) then entered the scene and pushed the climber up to the top of the hill. In the hindering condition, the scene was similar except that the second character (the hinderer) pushed the climber back down the hill and thus prevented the climber from achieving its goal. Afterward, children could choose or interact with the helper or the hinderer social agents. While initial findings yielded mixed results (*e.g.*, Hamlin et al., 2007, 2010; Holvoet et al., 2016; Margoni & Surian, 2018; Scarf et al., 2012), a recent large-scale and multi-lab replication study showed no preference for the helper over the hinderer social agents in children under 10 months (Lucca et al., 2025).

Three nonhuman species have also been tested on their prosocial preferences with the hill paradigm. Similar to children, dogs did not show any preference for helper or hinderer puppets (McAuliffe et al., 2019). Bottlenose dolphins (*Tursiops spp.*), tested with onscreen agents with different movements to those of hill-climbers to adapt to the socio-ecological aspects of dolphins, were able to discriminate pro- and anti-social behaviours and to predict preferential associations with the prosocial animated agent (Johnson et al., 2018). By contrast, bonobos (*Pan paniscus*) preferred hinderer over helper animated agents (Krupenye & Hare, 2018). The bonobos’ results contrast with the prosocial preference found in other nonhuman primates in other paradigms with human experimenters (Anderson et al., 2013, 2016; Brügger et al., 2021; Kawai et al., 2014; Subiaul et al., 2008). Note that bonobos did not show any preference for prosocial or antisocial agents in previous studies using different paradigms (Herrmann et al., 2013; Russell et al., 2008).

While this contrasting result could be due to an effect of the paradigm used (onscreen animated agents), the same bonobos replicated their preference for hinderers over helpers in subsequent experiments with human social agents (experiments 2 & 3 of Krupenye & Hare, 2018). The authors suggested that a preference for dominant-like behaviours might explain bonobos’ preference for hinderers. The movements of the helper and hinderer animated agents could be perceived as dominant- and subordinate-like behaviours. In an additional experiment, bonobos preferred animated agents showing dominant-like actions to ones showing subordinate-like actions (experiment 4 of Krupenye & Hare, 2018). Dominant individuals represent valuable social partners in this species, and it might be beneficial to interact with them preferentially; although, this is also true in other nonhuman primate species.

In this study, we tested another primate species sharing some behavioural and social traits with bonobos using the hill paradigm to see whether they would mirror the bonobos’ preference for hinderers. Like bonobos, Tonkean macaques (*Macaca tonkeana*) live in multi-male, multi-female groups, are highly tolerant and show prosocial interactions (Riley, 2010; Thierry, 2000; Thierry et al., 1994, 2004). They have demonstrated to track third-party interactions (Whitehouse & Meunier, 2020) and to understand goal-directed actions and attentional states (Canteloup et al., 2016, 2017; Canteloup & Meunier, 2017). For comparative purposes, we used the same experimental procedure used by Krupenye & Hare (2018) with the bonobos. Tonkean macaques could observe videos with the animated agents (*i.e.*, geometrical coloured shapes with googly eyes) several times. We then measured whether they would prefer a paper cutout representing the helper over one representing the hinderer (or vice versa). A non-differential rewarding procedure (*i.e.*, subjects received the same food outcome regardless of their choices) avoided shaping subjects’ preferences. A control condition was also conducted to check for potential effects of perceptual rather than social features on the videos. As Tonkean macaques have never been tested on social evaluation of prosocial behaviours, we considered that a helper, a hinderer, or no preference were equally likely, and therefore, made no predictions in this respect.

## Methods

### Subjects & testing conditions

Twelve Tonkean macaques (five males; 7.6 ± 6.3 years old) participated in the experiment voluntarily (Table S1). All subjects were captive-born, mother-raised, and came from one social group of 29 individuals (ten males; from 6 months to 28 years old) living in a wooded outdoor enclosure of 3 788 m^2^ with permanent access to an indoor room of 20 m^2^ at the Centre de Primatologie – Silabe de l’Université de Strasbourg (France). This group was fed once per day with dry pellets and once per week with fruits and vegetables and had access to water *ad libitum*. The data were collected in September 2023. Subjects were tested individually in a tunnel situated in a shelter directly accessible from their outdoor enclosure (video 1), inside which they were used to participate in touch screen cognitive experiments (Ballesta et al., 2020; Ballesta & Meunier, 2023; Fizet et al., 2017). A table was positioned at the end of the tunnel for the experimenter to place the video display, the rewards, and the paper cutouts representing the agents for the subjects to choose. All individuals in the group could participate in the experiment with the condition of being comfortable in the tunnel and with the experimental equipment. According to the STRANGE framework (Webster & Rutz, 2020), our selection protocol may have then biased our pool of subjects towards dominants, bold, highly motivated, and experienced individuals to experimental testing in the tunnel and/or with human experimenters. Before the experiment, a short familiarization step was conducted to verify whether each subject was comfortable entering the tunnel and watching videos on the screen (more details in the supplementary information). Five subjects have been tested before with two experiments on the social evaluation of human actors’ skills (Hirel et al., 2024; Hirel et al., in prep).

### Procedure & Design

The videos and animated agents in this study were the ones used by Krupenye et al. (2018) to test the bonobos. The animated agents were geometrically coloured shapes with two googly eyes, white sclera, and dark pupils, and they exhibited goal-directed movements. To control for any individual preferences, four different pairs of agents were used: red square and blue triangle, blue square and red triangle, orange pentagon and aqua trapezoid, aqua pentagon and orange trapezoid. Each subject watched different agent pairs in the test and the control session that did not share any physical characteristics (*e.g.*, red square and blue triangle during the test session, orange pentagon and aqua trapezoid during the control session). Both animated agents appeared the same amount of time on the screen and in contact with the third agent (the climber; see below). The types and roles of each agent (test: helper, hinderer; control: upward, downward) were assigned pseudo-randomly and counterbalanced across subjects (Table S1 & S2) and remained constant throughout the experiment. The initial arbitrary preference for one of the pair of shapes used in the test and control session was assessed on a separate day before the experiment, with no group preference found for a particular shape or colour (Table S3). The experiment consisted of a test and a control session. All the subjects participated in both sessions, with one session conducted per day and the presentation order counterbalanced between subjects.

#### Test session

First, the videos were presented to the subjects so they could observe the helping or hindering actions of the animated agents. In the test videos, the climber (a circle) entered the screen from below and repeatedly moved up on a hill, such as attempting to climb the hill but without success. After the third attempt, another agent entered the scene. In the helper video, the helper entered from below, pushed the climber up to the top of the hill and then returned down the hill to exit the screen (Figure 1a). In the hinderer video, the hinderer entered from above, pushed the climber back down the hill and then returned to the top to exit the screen (Figure 1b). Each subject watched the helper and hinderer videos four times, alternating in a loop. The type of the first video presented was counterbalanced between subjects. To control for the fact that the helper agent entered from the opposite side of the scene as the hinderer agent, all videos had a mirrored version. Each subject had half of the videos with the original version (*i.e.*, the helper entering from the right side, the hinderer from the left side) and half with the mirrored version (*i.e.*, the helper entering from the left side, the hinderer from the right side). The order of the original and mirrored versions was counterbalanced within and between subjects. The experimenter played the video loop only when the subject was attentive and looking toward the screen; otherwise, the experimenter tried to get the subject’s attention back by calling the subject and/or showing food. When the video loop ended, the experimenter removed the screen and gave a grape to the subject.

**Figure 1:**
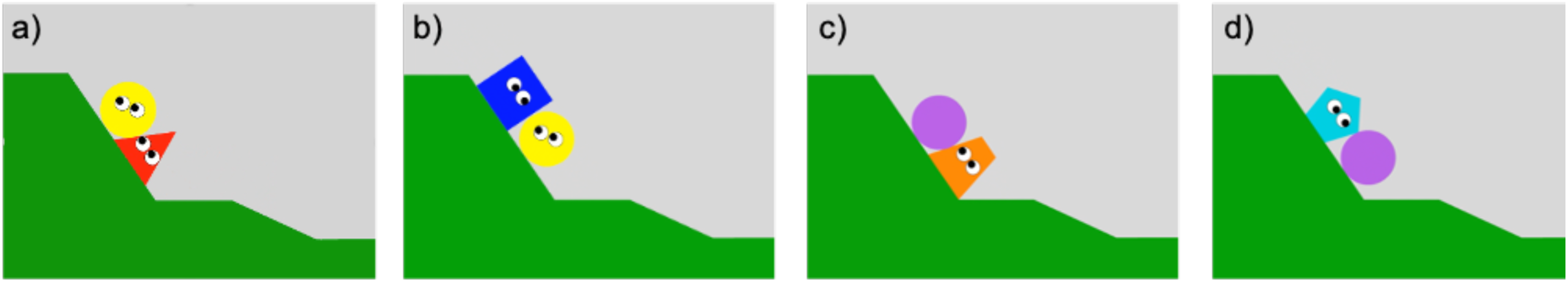
Frames from the video stimuli illustrating the a) the helper condition, b) the hinderer condition, c) the upward condition, d) the downward condition.

In the second part of the session, subjects could choose between two paper cutouts representing the helper and hinderer for four trials (video 1). Each trial started with the subject watching both helper and hinderer videos again once (the order of presentation and the version (original or mirrored) of the videos being counterbalanced within and between subjects). Then, the experimenter removed the screen, placed simultaneously two same-sized food items (banana chips) a few centimetres apart in the middle of the table, and covered them with the paper cutouts. The experimenter moved the two options to the opposite sides of the table so the subject could choose one. Pointing toward one of the options was used as a measure of preference. The experimenter placed the paper cutout selected next to the food, gave the food under it to the subject, and then took back both paper cutouts and the unchosen food reward. If no choice within 30 s or no clear pointing was made, the experimenter pulled the two options back and waited a few seconds before presenting the options again. If no choice was made in three consecutive trials, the experimenter stopped the session (which never happened). The location of the helper and hinderer paper cutouts on the table was counterbalanced and pseudo-randomized across trials.

#### Control session

The control session aimed to verify whether perceptual rather than social features, such as the movements of the animated shapes, could explain the subjects’ helper or hinderer preference. The procedure of the control session was similar to the test session except for the videos displayed to the subjects. Each control video was a variant of the test videos, with the climber replaced by an inanimate circle without eyes exhibiting no independent movement or goal-directed actions. The upward agent pushed the eyeless inanimate circle up the hill to control for the upward movement of the helper agent (Figure 1c). In contrast, the downward agent pushed the eyeless inanimate circle down the hill to control for the downward movement of the hinderer agent (Figure 1d). A preference for the upward (or downward) agent in the control session could explain a preference for the helper (or hinderer) agent in the test session by a sensitivity to the upward (or downward) movements of the shapes rather than their social interactions. The presentation order of the type (upward, downward) and the version (original, mirrored) of the videos were again counterbalanced between and within subjects.

### Data coding

Subjects’ choices were scored live by the experimenter. All sessions were videotaped with a GoPro9 camera and coded frame by frame by two observers, including one who was unaware of the study design and hypothesis, using Behavioral Observation Research Interactive Software (BORIS v.8.20; Friard & Gamba, 2016). For each session, videos were coded for a) the choices of the subjects toward one paper shape agent (inter-coder reliability: Cohen’s kappa, 𝜅 = 1, *N* = 96) and b) the duration of looking at the screen when the videos were displayed in the first step of each session and before each choice trial (inter-coder reliability: *ICC* = 0.819, *N* = 96). Inter-coder reliability was calculated using Cohen’s kappa coefficient for the choices and using Intraclass Correlation Coefficients (ICC; Koo & Li, 2016) for the duration of looking based on a single rating (*k* = 2 raters), two-way mixed-effects model and consistency. We calculated ICCs with the function icc from the package irr (version 0.84.1) in R (version 4.3.2; R Core Team, 2022).

### Data analyses

We assessed whether subjects preferred one animated agent over the other at the test (helper or hinderer) or the control session (upward or downward). We fitted a Generalized Linear Mixed Model (GLMM; Baayen, 2008) in R (version 4.3.2; R Core Team, 2022) with logit link function (McCullagh & Nelder, 1989) and a Binomial error structure using the function glmer of the package lme4 (version 1.1-35.1; Bates et al., 2015). In R, such an analysis of proportions is possible by using a two-column matrix with the number of successes and failures per individual as the response (Baayen, 2008). The response variable was a matrix comprising the number of choices for the helper and hinderer at the test session and for the upward and downward at the control session for each subject. The sample included 24 observations (session) from 12 subjects. This model included session (test, control) as the main predictor. To control for their potential effects, we also included sex and age of the subjects, order of presentation (whether test or control was presented first), and subjects’ attention to the videos. As subjects needed to observe the videos to obtain information about the animated agents’ actions, their amount of attention could have an effect on their choices at the trials afterwards. Subjects’ attention included in the model was the looking time per subject to the videos before the first trial (including the loop of videos and the two additional videos before the first trial) for each phase.

To account for individual differences, avoid overconfident model estimates and keep the type I error rate at the nominal level of 5 %, subject ID was included as a random intercept effect and session as a random slope within subject ID (Barr et al., 2013; Schielzeth & Forstmeier, 2009). Correlations between the random intercept and slope were not included in the model. Before fitting the model, session was dummy-coded and then centred. At the same time, age and subjects’ attention were z-transformed to a mean of zero and a standard deviation of one to ease the interpretation of the model estimates (Schielzeth, 2010) and aid model convergence. To test the effect of individual fixed effects, we conducted likelihood ratio tests (Dobson, 2018) that compared the full models with reduced models, each lacking one fixed effect at a time (Barr et al., 2013). We assessed whether collinearity was an issue using Variance Inflation Factors (VIF; Field, 2005), determined for a linear model using the function vif of the package car (version 3.1-2; Fox & Weisberg, 2011). We estimated the stability of the model by dropping the subjects one at a time from the data and comparing the estimates derived for models fitted to these subsets with those obtained for the full data set. We assessed overdispersion using a function provided by Roger Mundry (Mundry, 2023). The model revealed to be of good stability, was mildly underdispersed (dispersion parameter: 0.678, which may result in too conservative tests), and had no obvious issue of collinearity (maximum VIF: 2.3). We obtained confidence intervals of model estimates and fitted values using a parametric bootstrap (N = 1000 bootstraps; function bootMer of the package lme4).

## Results

In the test session, subjects chose the helper agent 26 times and the hinderer agent 22 times out of the 48 trials (Table S1). In the control session, subjects chose the upward agent 21 times and the downward agent 27 times out of the 48 trials (Table S2). The probabilities of selecting the helper in the test session or the upward agent in the control session were not significantly different from chance (Figure 2) and were not different from each other (*p* = 0.255; Table 1). The model revealed no significant effects of subjects’ attention (*p* = 0.562), sex (*p* = 0.808), age (*p* = 0.760), or the presentation order of the sessions (*p* = 0.380; Table 1). Subjects looked on average 37.2 ± 8.6 s (40 %) at the videos before the first trial in the test session (individual range: 24.6 s to 53.6 s) and 23.2 ± 8.2 s (39.3 %) in the control session (individual range: 13.9 s to 37.6 s).

**Figure 2:**
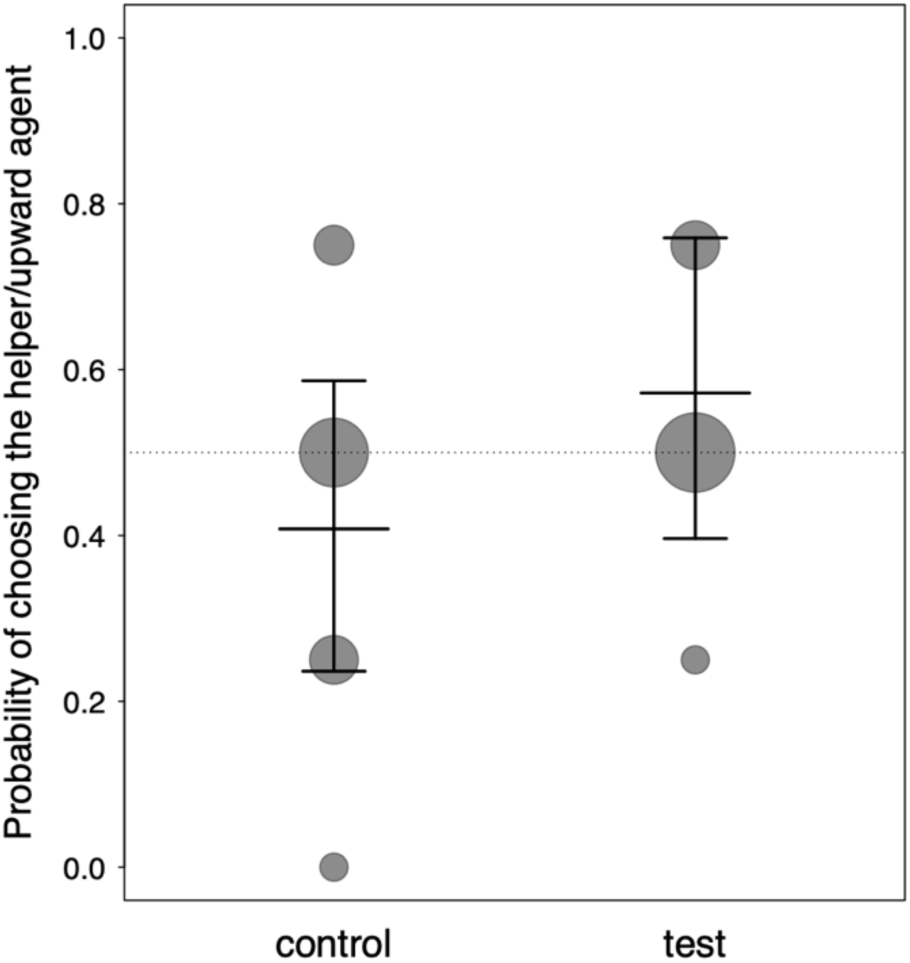
Probability of the subjects to choose the upward agent in the control session and the helper agent in the test session. The fitted model and its 95 % confidence limits are shown for both sessions. Data points depict the mean probability of choosing the helper/upward agent for each subject; the area of the points is proportional to the number of subjects (range: 1 to 8).

**Table 1:**
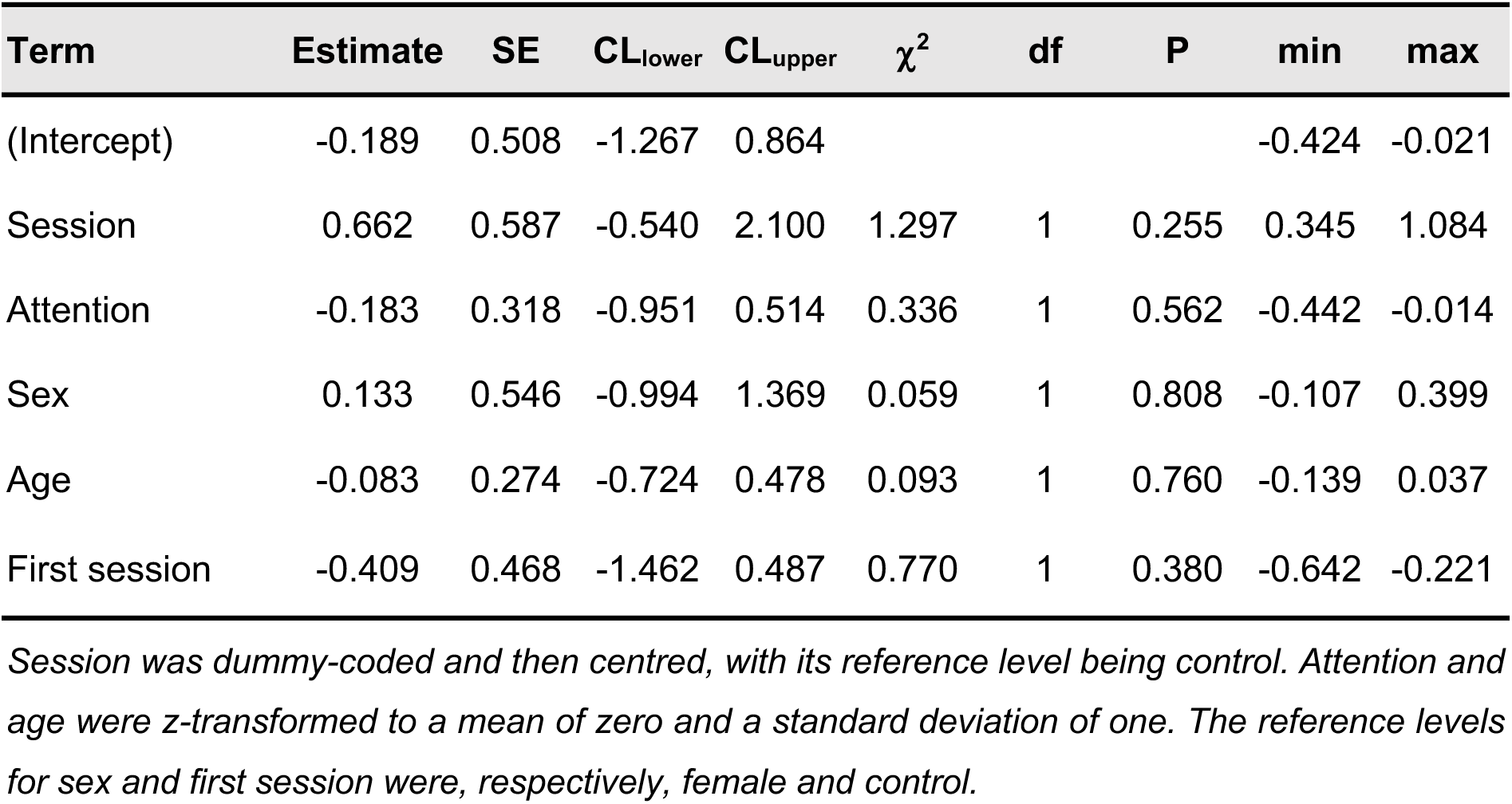
Results of the full model for the subjects’ choices in the test and control session (estimates together with standard errors, 95 % confidence limits, significance tests and the estimates range obtained when dropping levels of grouping factors one at a time).

| Term | Estimate | SE | CL<sub>lower</sub> | CL<sub>upper</sub> | $\chi^2$ | df | P | min | max |
| --- | --- | --- | --- | --- | --- | --- | --- | --- | --- |
| (Intercept) | -0.189 | 0.508 | -1.267 | 0.864 |  |  |  | -0.424 | -0.021 |
| Session | 0.662 | 0.587 | -0.540 | 2.100 | 1.297 | 1 | 0.255 | 0.345 | 1.084 |
| Attention | -0.183 | 0.318 | -0.951 | 0.514 | 0.336 | 1 | 0.562 | -0.442 | -0.014 |
| Sex | 0.133 | 0.546 | -0.994 | 1.369 | 0.059 | 1 | 0.808 | -0.107 | 0.399 |
| Age | -0.083 | 0.274 | -0.724 | 0.478 | 0.093 | 1 | 0.760 | -0.139 | 0.037 |
| First session | -0.409 | 0.468 | -1.462 | 0.487 | 0.770 | 1 | 0.380 | -0.642 | -0.221 |

*Session was dummy-coded and then centred, with its reference level being control. Attention and age were z-transformed to a mean of zero and a standard deviation of one. The reference levels for sex and first session were, respectively, female and control.*

## Discussion

Tonkean macaques did not prefer the helper nor the hinderer animated agent at the hill paradigm. The random choices found during the control session also indicated no effect of perceptual features of the shapes’ movements on the videos. Our results are similar to the ones found in children (Lucca et al., 2025) and dogs (McAuliffe et al., 2019) but contrast with the hinderer preference found in bonobos (Krupenye & Hare, 2018). However, only adult bonobos and not subadults preferred the hinderer. Although we managed to test Tonkean macaques of different ages (from one to 24 years old), we found no age effect in our study either. Therefore, Tonkean macaques did not show the same pattern of choices in the hill paradigm as the bonobos. While these two species share some behavioural and social characteristics (*e.g.*, high social tolerance, prosocial behaviours), it is still unclear whether Tonkean macaques monitor and keep track of others’ prosocial behaviours to choose their social partners or whether they show preferential interactions with dominants as much as bonobos do (Vervafcke et al., 2000; Yamamoto, 2015).

However, the testing conditions of the hill paradigm might not have motivated Tonkean macaques to pay attention to the videos and to evaluate the animated agents’ actions. Although the amount of attention our subjects paid did not affect their choices (similarly to children’s results; Lucca et al., 2025), overall, they looked at the videos not more than 40 % of the time during the test and the control session. The attention to the videos was limited, although the experimenter called the subject and/or showed food when the subjects looked away to try to attract their attention back. The amount of attention the bonobos paid to the videos would have been useful information for comparing differences in attention and their potential effect on the choices between the two species. The low level of attention of the Tonkean macaques could explain our findings, but above all, it indicates a lack of interest in the experimental stimuli. The actions of the animated agents were of no direct relevance to the subjects, and no food was involved in the interactions between the social agents. In contrast, previous studies showing a prosocial preference in nonhuman primates included social interactions between social agents that involved food (Anderson et al., 2013, 2016; Herrmann et al., 2013; Kawai et al., 2014; Russell et al., 2008; Subiaul et al., 2008). Whether nonhuman primates perceive others only as social tools to access resources or whether they also evaluate (positively or negatively) the prosocial tendencies of their conspecifics needs to be further investigated in future studies.

One potential limitation of our study is using videos and artificial animated shapes as social agents. Our subjects are highly habituated to screens as they have participated in touch screen cognitive experiments daily for several years (Ballesta et al., 2020; Ballesta & Meunier, 2023; Fizet et al., 2017). However, they may not have perceived the animated shapes as social agents. While primate species can recognize the content of pictures, how they process images and video stimuli is still unclear (Fagot et al., 2010). Great apes and Japanese macaques have been shown to perceive animated shapes as agents with goal-directed movements (Atsumi et al., 2017; Uller, 2004). Infant Japanese macaques, however, did not distinguish between objects in motion with or without eyes added (Tsutsumi et al., 2012). Tonkean macaques showed an understanding of goal-directed actions and attentional states of human experimenters (Canteloup et al., 2015; Canteloup & Meunier, 2017). However, their understanding of attentional states was based on rough cues such as body and head position rather than eye cues. Similarly, Tonkean macaques did not follow the eye gaze of a human experimenter in another study (Itakura, 1996). Therefore, our study’s use of googly eyes added on animated shapes with motion might not have been sufficient for Tonkean macaques to perceive them as social agents. In addition, Sliwa & Freiwald (2017) found a brain network that was exclusively activated when macaques observed videos of social interactions and not during the observation of object-object interactions or individuals/objects without motion. This suggests that macaques’ perception of the interaction between animated shapes or between real social agents might be different. In conclusion, Tonkean macaques did not prefer a helper over a hinderer animated agent (or vice versa). However, the artificial testing conditions might represent a limitation of the hill paradigm to investigate a prosocial preference in this species. In addition, the recent large-scale multi-lab replication study showing no prosocial preference of young infants in the hill paradigm calls into question the relevance of this paradigm for investigating social evaluation abilities in children too (Lucca et al., 2025). The hinderer preference of bonobos in the hill paradigm and replicated with other paradigms using human experimenters (Krupenye & Hare, 2018) remains puzzling and encourages further investigation. Specifically, we encourage further studies on the social evaluation of prosocial behaviours in Tonkean macaques and other species with paradigms using conspecifics or human actors as social agents. Additionally, in future studies, nonhuman primates’ prosocial behaviours and dominant-bias interactions need to be investigated to understand whether socio-ecological factors could explain the hinderer preference found in bonobos.

## Ethics statement

This study respects the European ethical standards and regulations (Directive 2010/63/UE). It has been approved by the internal ethical committee of the Centre de Primatologie – Silabe de l’Université de Strasbourg (SBEA 2022-03, registration n° B6732636). We only used positive reinforcement, and all individuals participated voluntarily. They had free access to the testing tunnel and could enter and leave as they wished. Their daily feeding regime was not affected by this experiment and water was available *ad libitum* at all times.

## Data accessibility

The videos, data, and statistical code associated with this article can be found at: https://osf.io/qyngx/

## Acknowledgements

We are grateful to the management, researchers, and animal keepers of the Centre de Primatologie – Silabe de l’Université de Strasbourg (https://www.silabe.com) for their approval and valuable support in conducting this study. We would like to thank Shadab Javanmard Paghaleh, Michele Marziliano, and Mathieu Legrand for their crucial help with the data collection and Nadja Vögtle for video coding. We are deeply grateful to Christopher Krupenye for generously sharing the experimental videos used previously with the bonobos.

## Author contributions

**Hirel Marie**: Conceptualization, Data curation, Formal analysis, Investigation, Methodology, Project administration, Visualisation, Writing – original draft, Writing – review & editing. **Meunier Hélène**: Resources, Writing – review & editing. **Rakoczy Hannes**: Funding acquisition, Writing – review & editing. **Fischer Julia**: Funding acquisition, Writing – review & editing. **Keupp Stefanie**: Conceptualization, Project administration, Supervision, Writing – review & editing.

